# ArMo: An Articulated Mesh Approach for Mouse 3D Reconstruction

**DOI:** 10.1101/2023.02.17.526719

**Authors:** James P. Bohnslav, Mohammed Abdal Monium Osman, Akshay Jaggi, Sofia Soares, Caleb Weinreb, Sandeep Robert Datta, Christopher D. Harvey

**Author notes:** These authors contributed equally to this work (James P. Bohnslav, Mohammed Abdal Monium Osman, Akshay Jaggi).

## Abstract

Characterizing animal behavior requires methods to distill 3D movements from video data. Though keypoint tracking has emerged as a widely used solution to this problem, it only provides a limited view of pose, reducing the body of an animal to a sparse set of experimenter-defined points. To more completely capture 3D pose, recent studies have fit 3D mesh models to subjects in image and video data. However, despite the importance of mice as a model organism in neuroscience research, these methods have not been applied to the 3D reconstruction of mouse behavior. Here, we present ArMo, an articulated mesh model of the laboratory mouse, and demonstrate its application to multi-camera recordings of head-fixed mice running on a spherical treadmill. Using an end-to-end gradient based optimization procedure, we fit the shape and pose of a dense 3D mouse model to data-derived keypoint and point cloud observations. The resulting reconstructions capture the shape of the animal’s surface while compactly summarizing its movements as a time series of 3D skeletal joint angles. ArMo therefore provides a novel alternative to the sparse representations of pose more commonly used in neuroscience research.

## 1 Introduction

Understanding how the brain produces behavior is a primary goal of neuroscience (Krakauer et al. 2017; Datta et al. 2019), and accomplishing this goal could yield insights that inform research in AI, robotics, and related fields. Given that behavior is ultimately expressed through the 3D movements of an animal’s body, methods for automatically extracting 3D pose from video recordings are essential for its study (Marshall et al. 2022). Moreover, since a great deal of neuroscience research involves mice, methods for measuring mouse movement kinematics are of particular interest.

In neuroscience research, the pose of an animal is often parameterized by the positions of defined locations (or “keypoints”) on the animal’s body, such as skeletal joints or superficial structures like the nose and ears (Mathis & Mathis 2020). Though keypoint tracking is highly informative, a limitation of this approach is that it reduces an animal’s body to a sparse set of points rather than capturing the body’s full shape. While this may suffice in animals with simple shapes or rigid exoskeletons, rodents have a highly deformable surface whose morphology is not fully captured by sparse representations.

Recent pose estimation studies of both human (Bogo et al. 2016; Huang et al. 2018) and animal (Zuffi et al. 2017; Badger et al. 2020) subjects have pioneered the use of articulated meshes as parametric body models. Articulated meshes are a commonly used class of 3D graphics models consisting of a polygon mesh and an underlying skeletal rig (e.g., Loper et al. 2015). Because these models represent the complete surface of an animal’s body, they can effectively capture the shape of structures that are poorly represented by a skeleton alone, such as fatty tissues. The parameters of an articulated mesh model can be fit to real data by minimizing the discrepancy between the shape and pose of the mesh and image-derived features such as keypoint positions and segmentation masks. These mesh fitting methods afford rich 3D reconstructions of a subject’s behavior from video recordings and could provide useful data for neuroscience research. However, there are currently no such methods for mice.

Here, we present ArMo, an articulated mesh model of the laboratory mouse that can be fit to keypoints and point clouds derived from multiview behavioral videos. We demonstrate ArMo by capturing the detailed body movements of head-fixed mice as they navigate in virtual reality (Harvey et al. 2009).

## 2 Related Work

### 2.1 Pose inference with articulated mesh models

Pipelines to reconstruct human shape and pose in 3D typically rely on a generative graphics model of the human body (Wang 2021). The most popular such model is SMPL, a linear model derived from thousands of high quality 3D scans of people in various poses (Loper et al. 2015). SMPL captures variation in pose using joint angles and variation in shape via a 3D-scan principal component representation. A variety of methods have used the SMPL model to estimate human pose and shape from images. SMPLify used pose priors and keypoint tracking to iteratively fit SMPL parameters to individuals in single images (Bogo et al. 2016). Further work extended this approach to multi-view video by incorporating temporal priors and a silhouette optimization term that relies on image segmentation (Huang et al. 2018).

Like approaches for humans, animal mesh reconstruction typically relies on parametric mesh models. The SMAL model is one such model for large quadrapeds such as cows, horses, and dogs that was learned using scans of animal figurines (Zuffi et al. 2017). Related work on 3D bird reconstruction circumvented the need for 3D scans by instead parameterizing shape variation using a set of bone lengths and an overall scale parameter (Badger et al. 2020). Like SMPL, both of these models are fit to real data using optimization procedures that incorporate keypoints, silhouettes, and priors on shape and pose.

### 2.2 3D pose inference in rodents

There has been a recent proliferation of methods for 3D pose inference in rodents (see Mathis & Mathis 2020 and Marshall et al. 2022 for a more indepth review). Approaches such as DeepLabCut 3D use neural networks to independently extract 2D keypoints from multiple view angles and then triangulate them to obtain 3D positions (Nath et al. 2019). Alternatives, such as DANNCE, instead project image features into 3D space and then use volumetric convolutions to regress 3D keypoint positions directly (Dunn et. al. 2021; Schneider et. al. 2022). To improve on these algorithms, some methods have leveraged model-based approaches. For example, GIMBAL instantiates spatiotemporal priors on the overall pose of the animal via a hierarchical von Mises-Fisher-Gaussian model, which results in better inference of 3D keypoint positions than naive triangulation (Zhang et al. 2021). Another approach called the ACM (anatomically constrained model) uses Kalman smoothing and pose priors to accurately reconstruct 3D skeletal kinematics in freely behaving rodents over a wide range of sizes (Monsees et al. 2022). Other methods have fit simplified body models to rodents. 3DDD Social Mouse Tracker, for example, uses a particle filtering algorithm to fit pairs of prolate spheroids to point cloud and keypoint observations of mice (Ebbesen & Froemke, 2022).

## 3 Approach

To develop an articulated mesh model of the laboratory mouse, we started with an artist generated 3D model and simplified it by removing the joints corresponding to digits on the forepaws and hindpaws. The resulting model consists of a skeleton with 30 joints, a mesh with 1,803 vertices and 3,602 faces, and an 1803 × 30 matrix of skinning weights that determines mesh shape as a function of joint angles (Fig. 1; Supplementary Video 1). To facilitate the fitting of the model to keypoint observations, we also manually specified which subsets of vertices on the mesh correspond to keypoints of interest.

**Fig. 1.**
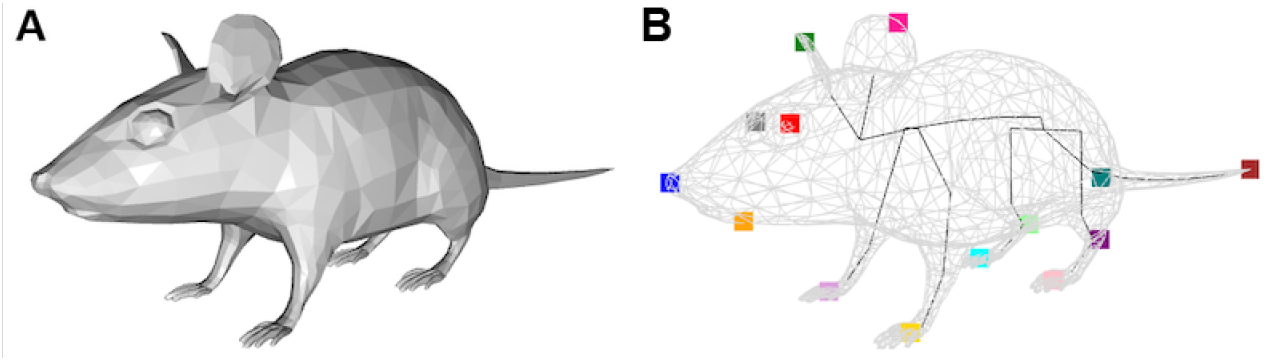
ArMo mesh model. **A**: Triangle mesh surface as it appears in the rest pose. **B**: Wireframe view of the mesh with the underlying skeleton in black. Colored points indicate keypoints used for fitting.

Knowledge of mouse anatomy was leveraged to simplify the shape and pose parameters. In particular, to reduce the number of trainable parameters, we used weight sharing to enforce left-right symmetry in the bone lengths. Similarly, we froze certain degrees of freedom in the joint angles to ensure a measure of anatomical accuracy in the mouse’s pose (for example, by preventing the knee from rotating laterally). After these constraints were applied, the model had 93 free parameters. It is important to note that although the modeled skeleton is made up of “joints” and “bones”, it should only be thought of as a useful parameterization of pose rather than an anatomically precise description of real joint angles and positions in the animal’s body.

To parameterize and skin the mesh model, we used the procedure introduced in Badger et al. 2020 (described fully in the next section). To fit the mesh model’s pose and shape to videos of real mice, we enumerated an objective function that quantifies how well a given mesh configuration matches data-derived keypoint and point cloud observations. As in other mesh optimization approaches, we leverage the fact that the loss function is differentiable with respect to the mesh parameters to fit the pose and shape via end-to-end gradient-based optimization.

### 3.1 Pose Parameterization and Skinning Algorithm

Let *J* = 30 and *N* = 1, 803 denote the number of skeletal joints and mesh vertices in the model, respectively. The *J* joints in the skeleton are hierarchically arranged in a kinematic tree specifying which joints are connected by bones. The root joint is indexed by *j* = 1, and its “parent-relative” descriptors are defined with respect to the global coordinate system. The mesh vertices are connected by 3,602 triangular faces, and their positions are determined by the rotations and positions of the joints via linear blend skinning.

The pose of the model is parameterized by four trainable variables: a set of bone lengths 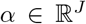, where *α_j_* is a scalar controlling the length of the bone connecting the *j*’th joint and its parent joint; a set of pose parameters 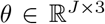, where *θ_j_* denotes the axis-angle representation of the *j*’th joint’s rotation in its parent’s reference frame; a translation vector 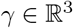 controlling the global position of the root node in the world; and a scale parameter 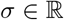 controlling the overall size of the mesh model. Note that for all *j*, we compose the rotation specified by *θ_j_* with the rest pose rotation for joint *j*. Therefore, these joint angle values are not absolute, but rather differences from the rest pose

Let *M*(*α, θ, γ, σ*) denote the blend skinning function that takes in the pose parameters and returns a mesh 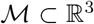 (note that while 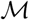 is a continuous surface defined by both its vertices and faces, we use the subscript notation 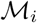 to index the *i*’th of its *N* discrete vertices). We chose to use linear blend skinning to determine the posed positions of the mesh vertices, but note that any other differentiable skinning function could also work in principle. The blend skinning parameters include: a zero pose 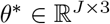 defining the parentrelative rotations of each joint at rest; a set of rest pose joint offsets 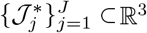 specifying each joint’s position in the reference frame defined by its parent joint; a set of rest pose vertex positions 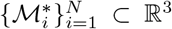; and a matrix of skinning weights 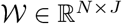. We assume that the root node is centered at the origin (i.e., 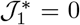) to ensure that the global translation parameter *γ* is not degenerate. Finally, for each of the *K* keypoints of interest, we define a subset of mesh vertices 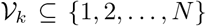. Each keypoint is assumed to lie at the centroid of its respective vertex-set.

In order to skin the model, we first determine the pose of the skeleton, which is defined by the global rotation and position (henceforth the “transform”) of each of its *J* joints. As is standard in skeletal rigging, we represent the global transform of joint *j* as a matrix 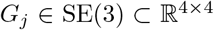, and to compute *G_j_*, we determine the local transforms associated with each of joint *j*’s ancestor nodes and propagate them down the skeletal tree. More formally,

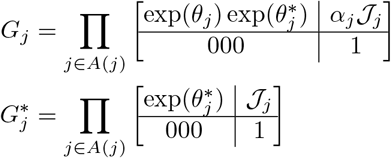

where *A*(*j*) represents the ordered set of joint *j*’s ancestors (starting with the root node and terminating with joint *j* itself), 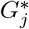 represents joint *j*’s global transform in the rest pose with the default bone lengths, *G_j_* represents the global transform with the effects of bone length scaling and posed joint rotation accounted for, and exp(·) denotes the Rodrigues formula, which is a differentiable transform that converts axis-angle representations of rotations into their associated 3 × 3 rotation matrices.

Next, to determine the positions of the mesh vertices, we use linear blend skinning and apply the scale and translation parameters:

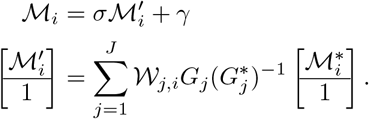

Finally, to determine the position of each mesh keypoint, we simply compute the mean position of its associated mesh vertices. More formally, let 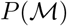 denote the function that takes in the mesh and returns a set of keypoint positions 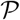, where 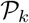 denotes the position of the *k*’th keypoint. Then

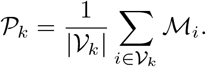

### 3.2 Pose Optimization

Using a procedure similar to SMPLify (Bogo et al. 2016) and 3D bird reconstruction (Badger et al. 2020), we fit the pose parameters to data by minimizing a loss function with two data terms and four regularization terms. The data terms capture how well the posed mesh corresponds to the keypoint and point cloud observations, and the regularization terms encode anatomical and temporal priors on the shape and pose.

To incorporate temporal priors, we simultaneously fit the pose across all *T* frames in a temporally contiguous segment of video. More formally, let 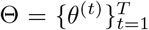 and 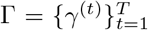 respectively denote the relative joint angles and translations for each timestep. Similarly, for convenience, let 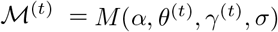 and 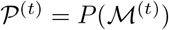 denote the mesh vertices and mesh keypoints associated with the pose at timestep *t*. Note that the shape parameters *α* and *σ* are shared across timesteps. The loss function *L* can be written as follows, with each *λ* coefficient specifying the non-negative scalar weight of its associated loss term:

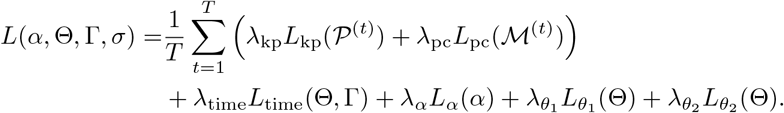

The keypoint loss *L*_kp_ measures the distance between the reprojections of the mesh keypoints onto each sensor and the corresponding image-derived keypoint predictions. It has the following functional form:

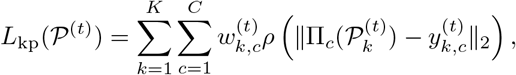

where *ρ* is the Geman-McClure function, which has been used in prior works for its robustness to noisy outliers (Geman 1987; Bogo et al. 2016; Badger et al. 2020), Π_c_ is the pinhole camera projection function mapping 3D points in world coordinates to the pixel space of camera 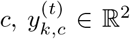 is the estimated pixel location of keypoint *k* in camera *c*’s image at time *t*, and 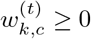 is the confidence associated with 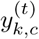 (Bogo et al. 2016).

The point cloud loss *L*_pc_ quantifies the overall shape mismatch between the point cloud and the mesh at a given timestep. Whereas prior work (e.g., Huang et al. 2018, Badger et al. 2020) has typically used 2D segmentation masks to provide shape supervision, here we leverage point clouds derived from depth imaging to incorporate 3D shape information more directly. To do so, the point cloud loss computes the sum of squared Euclidian distances from each point in the point cloud to the nearest face on the triangular mesh. This value can be approximated by sampling points from the surface of the mesh and computing pairwise distances. Because some regions of the animal’s surface may be absent in the point cloud due to occlusions, this distance term is unidirectional, unlike the more commonly used Chamfer distance. It has the following functional form:

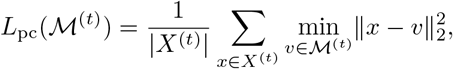

where 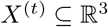 denotes the point cloud at time *t*.

The temporal loss *L*_time_ encourages smooth movements by penalizing the magnitude of the difference between the pose parameters at successive timesteps (Huang et al. 2017; Biggs et al. 2018). More formally,

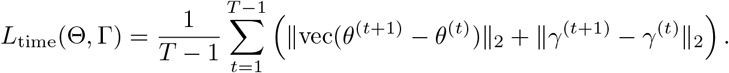

We also use three other regularization terms described in Badger et al. 2020. The bone loss term *L_α_* penalizes the model for assigning bone lengths outside of a predefined range of allowable values. It is defined as

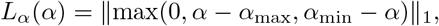

where max is applied element-wise and *α*_min_, 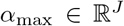 define the bounds outside of which a linear penalty is applied to each entry of *α*. Similarly, *L*_*θ*_1__ linearly penalizes joint rotations outside the bounds defined by *θ*_min_, 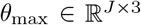:

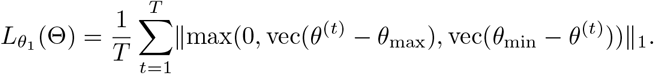

Finally,

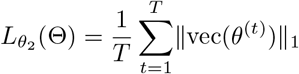

penalizes deviations from the rest pose.

We minimize the loss using gradient descent, with the optimization divided into two phases. In the first phase, we optimize *σ*, Γ, and 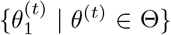 to determine the overall scale, position, and orientation of the mouse in global coordinates. In the second phase, we allow all of the parameters except for *σ* to freely vary, thereby learning *α* and the remaining joint angles in Θ.

In order to fit the mesh model to video data, we implemented this optimization procedure in PyTorch and leveraged the package’s auto-differentiation and hardware accelerator functionalities (Paszke et al. 2017). We also relied on a number of features from PyTorch3D, including its implementations of camera projections, differentiable sampling, and point cloud related functions (Ravi et al. 2020). To perform the optimization itself, we used an Adam optimizer (Zhang 2018) to fit the mesh in batches of 300 contiguous timesteps (corresponding to five seconds of 60 fps video). Simultaneously fitting the mesh for all the frames in a long recording is infeasible, yet the shape parameters for a single mouse should not vary over time. To get around this, we first fit both the shape and pose with no temporal regularization (i.e., by setting λ_time_ = 0) to a noncontiguous, randomly sampled subset of the frames. Using the shape (*α* and *σ*) learned from this random batch, we then fit the pose (Θ and Γ) for the remainder of the video, thereby disentangling shape and pose in our reconstruction of the behavior.

## 4 Experiments

### 4.1 Data acquisition

We built an experimental setup to capture the full-body movements of head-fixed mice running on a spherical treadmill (Fig. 2). To minimize occlusions and maximize the quality of the 3D reconstruction, we placed four stereo depth cameras (Intel Realsense D435) around the treadmill. Each camera has two closely-spaced infrared sensors (yielding a total of eight images per time point) and an infrared projector that illuminates the scene with a random dot pattern that aids depth estimation in less textured regions of the image, such as the side of the mouse. Videos were acquired at 640 × 480 resolution and 60 frames per second, and a very short shutter time of 750 microseconds was used to minimize motion blur. Using this setup, we recorded videos of mice as they performed a virtual navigation task. In addition, OpenCV (Bradski 2000) was used for camera calibration and stereo matching.

**Fig. 2.**
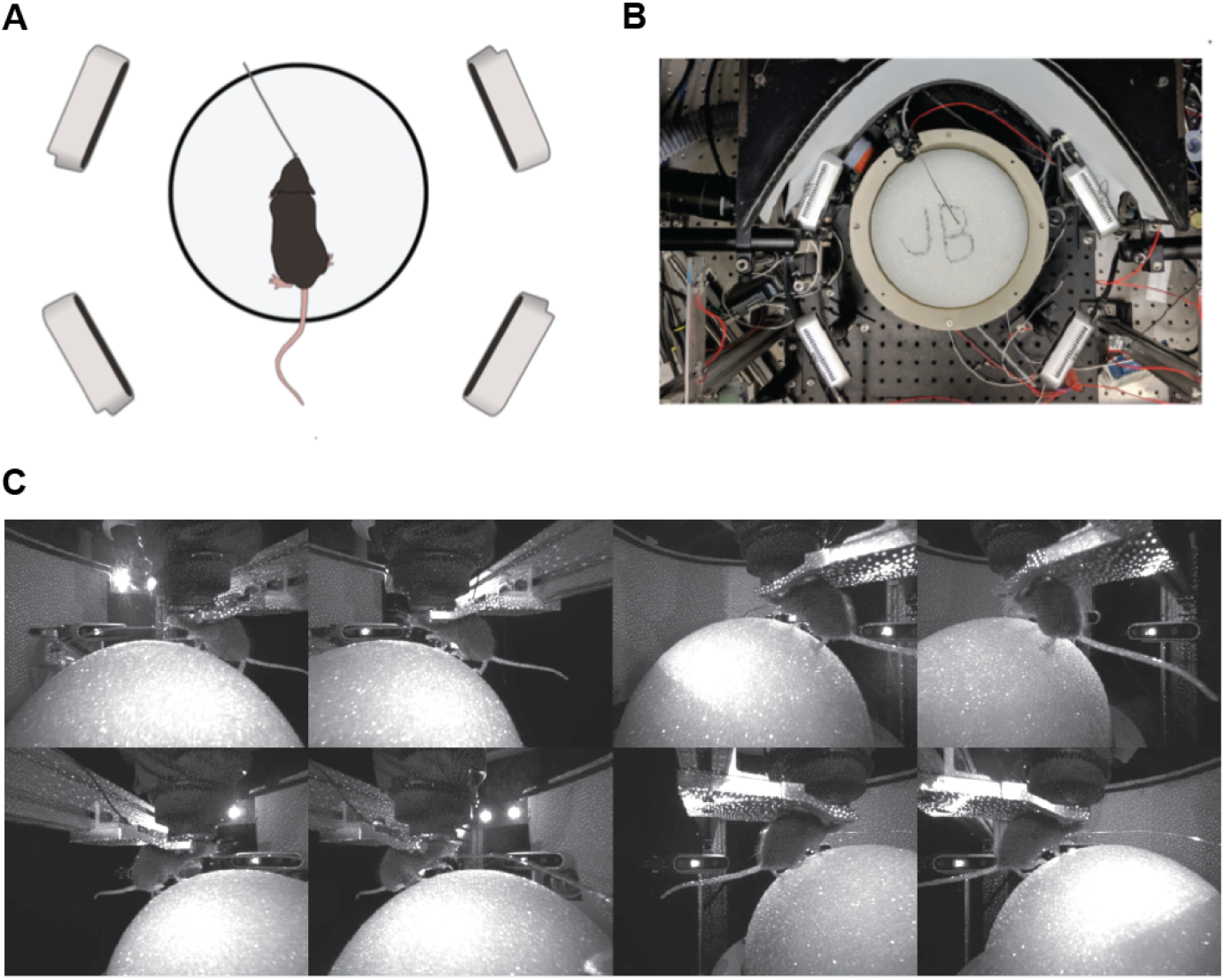
Behavioral setup and data acquisition. **A**: Recording rig, featuring a mouse on a spherical treadmill, a thin tube for delivery of water rewards, and four Intel RealSense D435 depth cameras. **B**: Photograph of the behavior setup corresponding to the schematic in (A). **C**: Example images from the infrared sensors.

### 4.2 Feature Extraction

We performed 2D keypoint localization and semantic segmentation simultaneously using a multi-headed deep convolutional neural network (Fig. 3A) with a pretrained HRNet backbone (Wang et al. 2020). The tracked keypoints included the nose, mouth, eyes, ears, forepaws, ankles, hindpaws, tail base, and tail tip. Semantic segmentation was used to detect the mouse, the spherical treadmill, and any occluders between the mouse and the camera (such as the hardware used for head restraint). The network performed well on validation data, with low pixel distance between real and estimated 2D keypoints (Fig. 3B) and high intersection over union with the true segmentation masks (Fig. 3C).

**Fig. 3.**
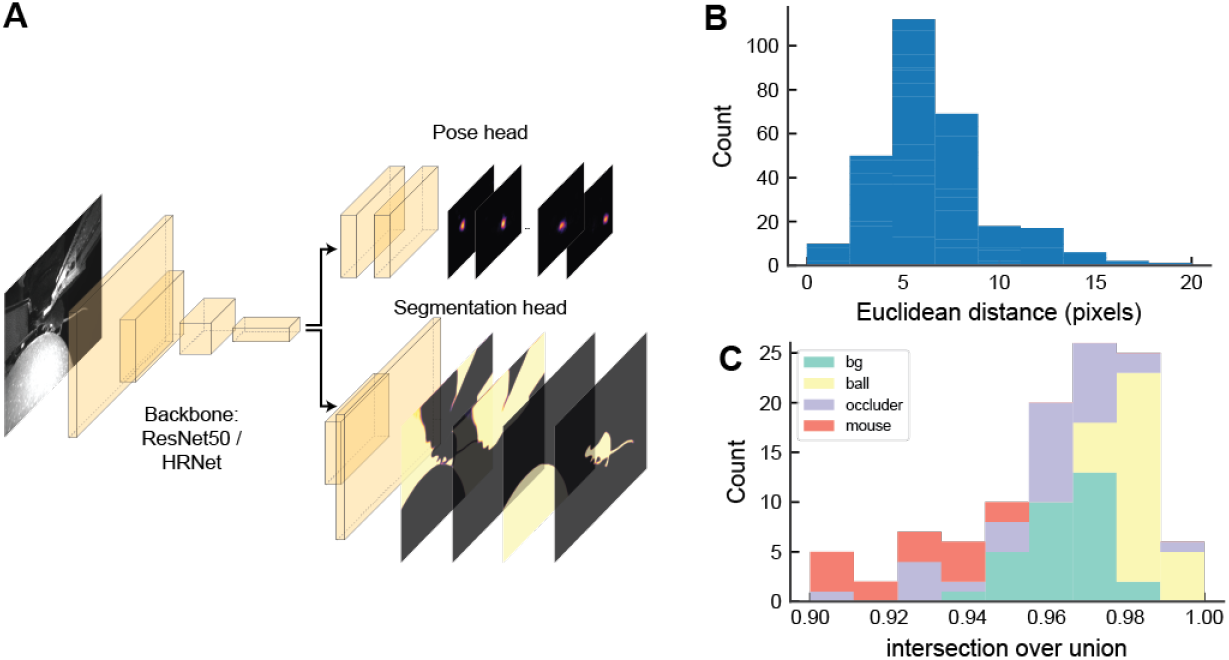
2D keypoint and segmentation network. **A**: Network architecture: An image is passed through a convolutional backbone which results in a feature map. This map is passed through two separate heads, one for 2D keypoint estimation and one for semantic segmentation. **B**: Distribution of distances between human-annotated and model-predicted 2D keypoint locations in the validation dataset. **C**: Distribution of intersection over union scores for human-annotated vs. model-predicted segmentation masks in the validation dataset.

To generate point clouds, we first performed offline stereo matching on the paired images from each camera, producing a map of estimated depth values for each pixel. The depth images were then filtered using the mouse segmentation masks and projected into a common 3D coordinate space. To refine the point cloud, we first cropped it to the 3D bounding box of the behavioral rig and then removed points whose average distance to their nearest neighbors exceeded an outlier threshold. This procedure resulted in reasonable looking point clouds that captured the shape of the mouse based on inspection of example frames (Fig. 4; Supplementary Video 2).

**Fig. 4.**
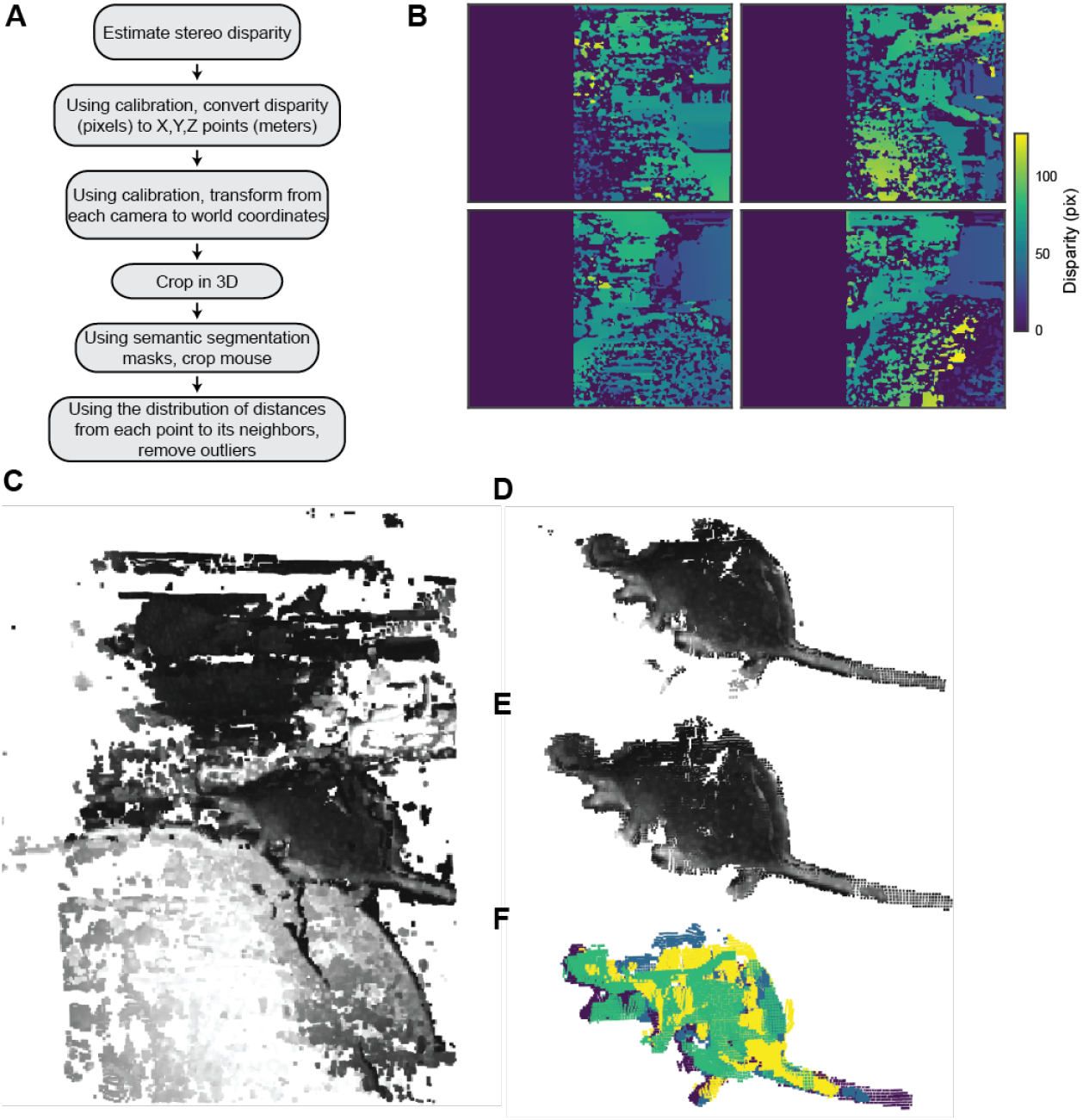
Point cloud pipeline. **A**: Overview of steps. **B**: Estimated depth maps for an example frame. The high noise-level is mostly attributable to reflections from the Styrofoam sphere that supports the mouse. **C**: Example point cloud in world coordinates, colored by pixel values in the original infrared images. **D**: Point cloud after masking with the estimated mouse silhouettes. **E**: Point cloud after cleaning with nearest neighbors. **F**: Same as (E), but colored by the identity of the camera from which the points originated.

### 4.3 Model Fitting

To test the optimization procedure, we fit ArMo to keypoint predictions and point cloud observations derived from three hours of the acquired video data (Fig. 5B; Supplementary Video 3). Visual inspection of the resulting mesh fit videos revealed that the procedure nicely recovered the shape of the mouse across views and captured its running on the spherical treadmill (Fig. 5A; Supplementary Video 4).

**Fig. 5.**
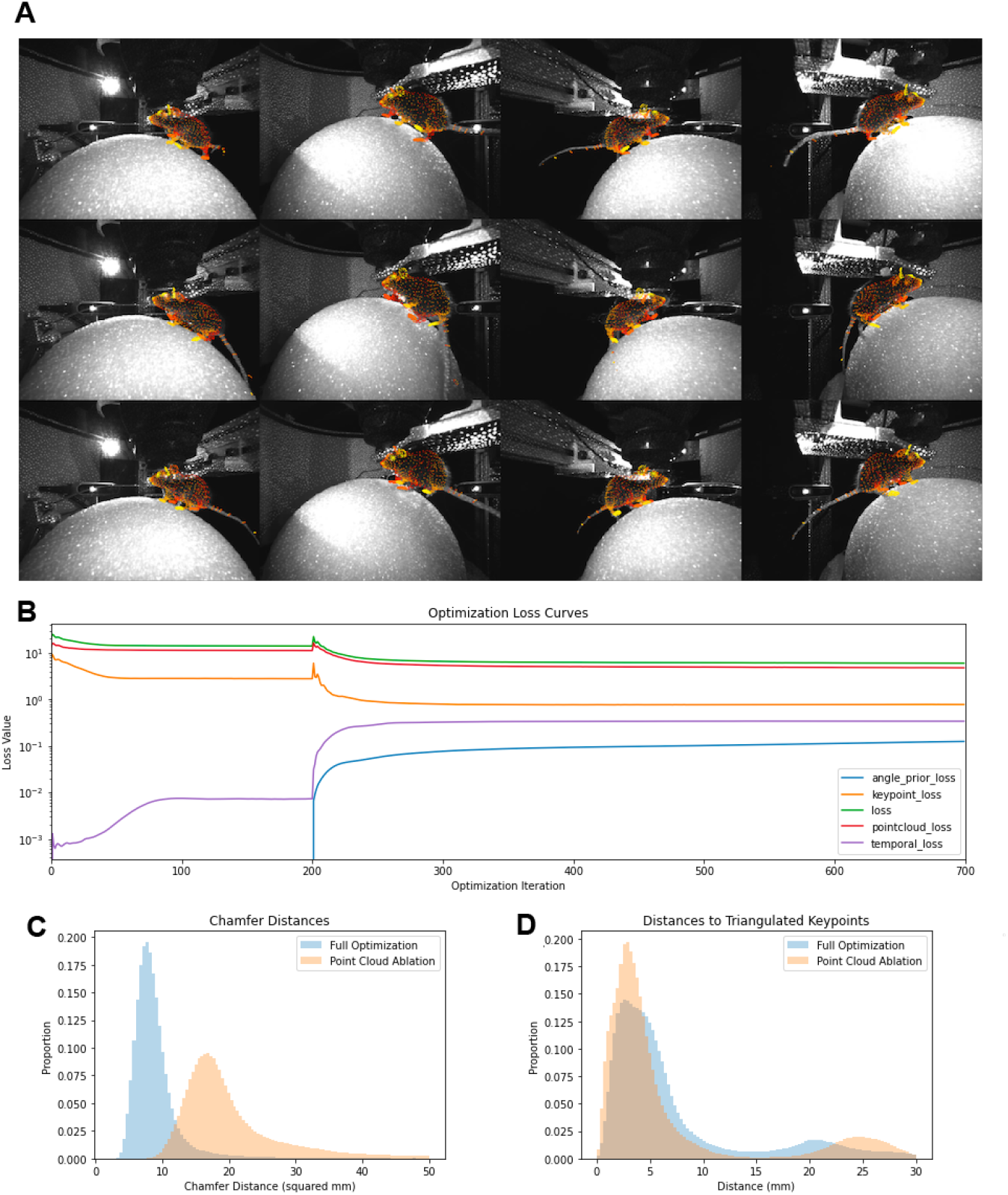
Model fitting. **A**: Mesh reconstructions overlaid on the original images. The color of each mesh vertex is shared across camera views. Each row corresponds to a timestep, and each column corresponds to a camera. **B**: Progression of each loss term during fitting. Note the log scale. **C**: Distributions of cloud-to-mesh distances across timesteps for the full optimization procedure (blue) and the point cloud ablation (orange). The cloud-to-mesh distance measures the average squared distance from each point cloud element to the mesh surface. Lower values indicate a greater shape similarity between the real and reconstructed mouse. **D**: Distributions of keypoint distances for the model fits shown in (C). Keypoint distance is computed as the Euclidean distance between a data-derived 3D keypoint estimate produced via direct triangulation and the corresponding point on the mesh.

We also performed an ablation experiment to determine how the point cloud data contributed to the mesh reconstruction. We found that the point cloud term improved the similarity between the mesh and the point cloud (Fig. 5C) while only negligibly increasing the median keypoint error (Fig. 5D). While this improvement was expected given that the optimization procedure directly minimizes this value, it nonetheless suggests that the keypoints alone do not fully specify the pose, and that additional sensor data (e.g., depth) can further constrain the remaining degrees of freedom.

To analyze the behavior of the mice, we performed principal component analysis on the 3D joint positions (excluding points in the tail) output by ArMo. Six principal components collectively explained 84.3% of the variance in the dataset (Fig. 6A). The components captured pose states characteristic of various behaviors, such as turning (component 1) and running (component 3) (Fig. 6C-D), and exhibited corresponding dynamics over time (e.g., smooth oscillatory dynamics in component 3 during running bouts; Fig. 6B).

**Fig. 6.**
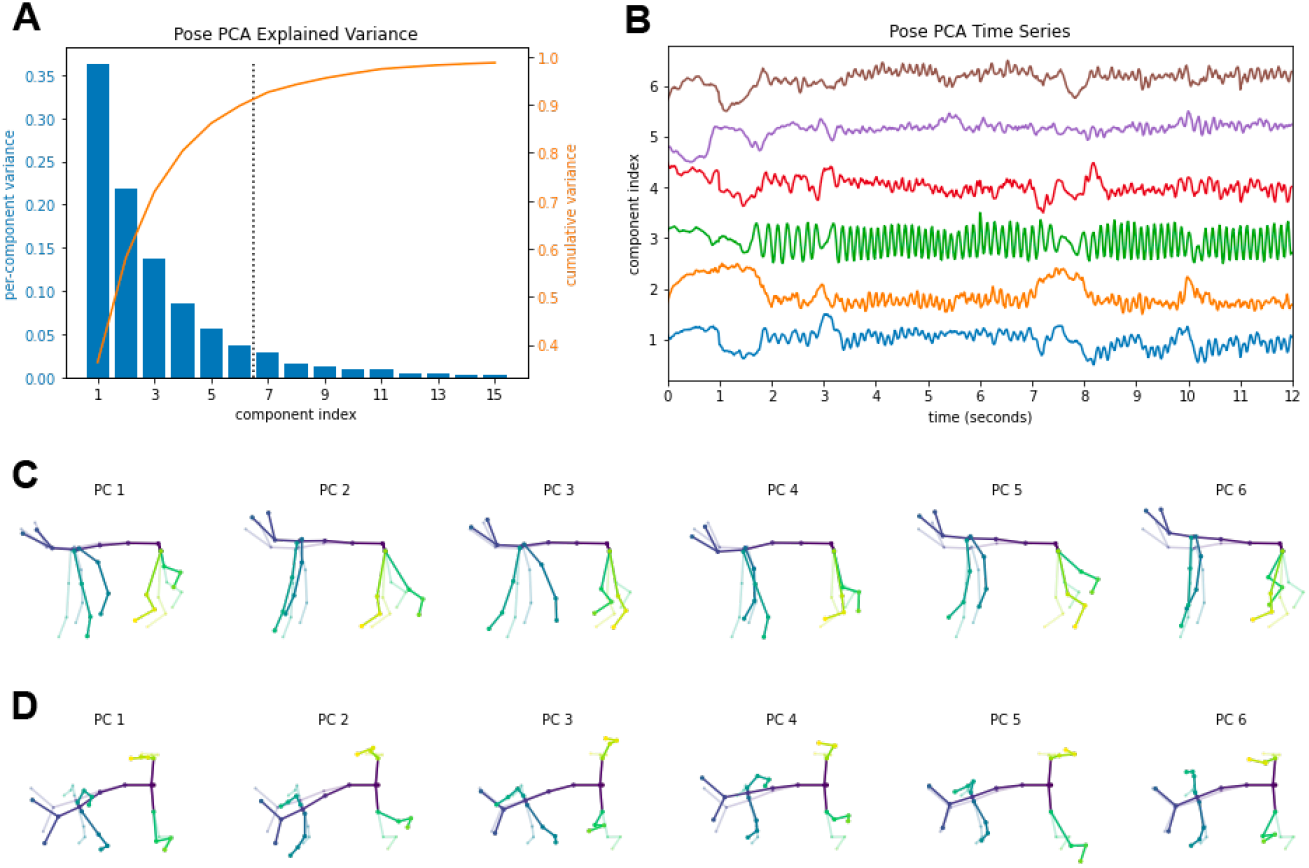
Principal component analysis (PCA) of ArMo pose states. **A**: Variance explained by each component individually (blue) and cumulatively (orange). Only the first six components (delimited by the dashed line) are shown in (B), (C), and (D). **B**: PCA projections for 12 seconds of example data, showing behavioral features such as high frequency oscillations in component 3 during running. **C**: Deviations from the mean pose in the direction of each component (side view). The solid lines indicate reprojections of the principle component axis into the 3D skeletal joint coordinate space, and the transluscent lines indicate the mean pose. **D**: Same as (C) for top-down view.

## 5 Conclusions

ArMo provides a novel alternative to 3D pose reconstruction in mice that captures the animal’s surface and movements more holistically than sparse representations like keypoints. To validate this method, we showed that it can reconstruct the movements of mice running on a spherical treadmill. We also leveraged point cloud data to aid mesh fitting. Future work could build on ArMo by generalizing this approach to freely moving mice, amortizing the fitting procedure by inferring mesh parameters directly from video (Kolotouros et al. 2019; Badger et al. 2020), or more deeply characterizing how mesh-based representations of pose differ (e.g., in their dynamics or dimensionality) from 3D keypoint representations.

## Acknowledgements

A.J. is supported by National Institute of General Medical Sciences grants T32GM007753 and T32GM144273. S.S. is supported by a European Molecular Biology Organization Postdoctoral Fellowship and a Brain & Behavior Research Foundation Young Investigator Grant. C.W. is a fellow of the Jane Coffin Childs Memorial Fund for Medical Research. S.R.D. is supported by NIH grants U19NS113201, RF1AG073625, R01NS114020, and U24NS109520, the Brain Research Foundation, and the Simons Collaboration on the Global Brain. C.D.H. is supported by NIH grants DP1 MH125776 and R01 NS089521. Portions of this research were conducted on the O2 High Performance Compute Cluster at Harvard Medical School. We thank our colleagues in the Harvey and Datta Labs for their feedback and support over the course of this project. The content is solely the responsibility of the authors and does not necessarily represent the official views of the National Institute of General Medical Sciences or the National Institutes of Health.

## Author Contributions

J.P.B. and C.D.H. conceived of the project. J.P.B. and C.D.H. developed the approach with contributions from A.J. and M.A.M.O. J.P.B. and S.S. built the hardware. S.S. collected the data. J.P.B. labeled the data. J.P.B., M.A.M.O., A.J., and C.W. wrote the software and analyzed the data. M.A.M.O., A.J., J.P.B., C.W., C.D.H., and S.R.D. wrote the manuscript with input from all authors.

## Notes

### Competing Interest Statement

The authors have declared no competing interest.

